# Optimising voyages for biodiversity: rerouting vessels around ocean giants can have minimal impact on shipping

**DOI:** 10.1101/2025.09.26.678754

**Authors:** Ryan R. Reisinger, Przemyslaw A. Grudniewski, Freya C. Womersley, David W. Sims, Adam J. Sobey

## Abstract

Ship strikes are a significant and growing threat to marine megafauna, yet few mitigation measures are implemented at scale due to perceived economic costs to shipping. Here, we present a proof of concept for integrating biodiversity considerations into commercial voyage optimisation, using priority aggregation sites for the endangered whale shark (*Rhincodon typus*) as a case study. We simulated eight port-to-port voyages for two vessel classes--a crude oil tanker and a container ship--under three routing scenarios: baseline optimisation, speed reduction to 10 kts within core habitats, and complete avoidance of these areas. Across routes, fuel-use changes ranged from −0.13% to 9.65%, with minimal impacts (<1%) for most long-distance voyages. Results indicate that speed reduction is the more efficient mitigation for short voyages, while area avoidance is preferable for longer voyages, with impacts varying by vessel type and operational constraints. Incorporating dynamic, species-specific habitat layers into voyage planning could enable targeted ship-strike mitigation with negligible disruption to global trade. Adoption of such measures – supported by improved data pipelines, real-time forecasting, and integration into regulatory and incentive frameworks – offers a scalable pathway to align biodiversity conservation with decarbonisation goals in the maritime sector.

## Introduction

World trade is enabled by a growing fleet of vessels carrying a range of produce all over the world. Presently, the global merchant fleet carries over 80% of goods (*1*) across 4,702 ports in 170 countries (*2*). In 2023, this global merchant fleet consisted of 105,500 vessels >100 gross tons, of which 56,500 were larger vessels, >1000 gross tons (*3*, *4*). The rate of increase in these vessels has been rapid, with only 1,441 vessels >100 gross tons in 1993 (*5*). This is expected to continue increasing, with world trade of goods expected to double by 2050 (*6*). This will increase the number of required ships by up to three times with the majority of these either being large ships or very large ships (*7*).

While shipping can be efficient, it is not without impact on ocean ecosystems, for example through noise pollution, chemical pollution, introduction of invasive species, marine litter, and the indirect effects of greenhouse gas emissions--global shipping accounted for ∼3% of anthropogenic CO_2_ emissions in 2018 (*8*). This impact will increase as the number of ships grows. To formulate routes that optimise both operational efficiency and safety, Voyage Optimisation Solutions (VOS) are widely utilised in commercial shipping. Studies have demonstrated reductions in carbon emissions and fuel costs ranging from 5% to 15% (*9*) and when combined with other Machine Learning approaches have the potential to save up to 18% (https://rina.org.uk/industry-news/naval-architecture/greater-than-the-sum-of-their-parts-merging-green-technologies/). State-of-the-art systems consider a comprehensive set of factors, including high-fidelity meteorological and oceanographic data, navigational restrictions, safety constraints, dual-fuel operations, variable departure times, high-fidelity vessel performance models and just-in-time arrival at ports. However, they will change the routes that vessels take across the oceans and currently there is no way to consider ocean biodiversity. Currently, physical collisions with wildlife (‘ship strike’) has been identified as a key conservation concern for various large marine vertebrates (hereafter ‘marine megafauna’) (*10*).

These collisions often physically injure or kill wildlife, may cause vessel damage, and pose risks to human safety (*10*, *11*). Ship strikes have been recorded for at least 75 marine species, including whales, dolphins, porpoises, dugongs, manatees, various sharks, seals, sea otters, sea turtles, penguins, and fish. However, reports of collisions with smaller species are rare, likely due to biases in reporting (*10*). The International Whaling Commission’s (IWC) Global Vessel Strikes Database includes 933 records of collisions between ships and cetaceans (whales, dolphins and porpoises) from 1820-2019 (*12*). However, collisions may go undetected or unreported, resulting in underestimates of the number of animals struck by vessels, besides various other data gaps and biases (*12*). For animals which sink when dead like the world’s largest fish, the endangered whale shark (*Rhincodon typus*), lethal strikes have been recorded through indirect measures, such as depth recording tags showing likely dead individuals sinking to the sea floor in busy shipping lanes (*13*). A study showed that from over 200 whale sharks fitted with data logging tags, 24% ceased transmitting data in heavily vessel trafficked areas, with lethal strikes suggested as being responsible for a substantial portion of these (*13*). Collisions may represent considerable, unquantified mortality to populations. For example, it has been estimated that in southern California around 23 blue, fin and humpback whales were killed by ship strikes in summer each year from 2012-2018 (*14*). For blue whales, this exceeds the United States Marine Mammal Protection Act’s (https://www.fisheries.noaa.gov/topic/laws-policies/marine-mammal-protection-act) Potential Biological Removal—the “maximum number of animals, not including natural deaths, that may be removed from a marine mammal stock (i.e., population) while allowing the population to maintain or recover to its optimum sustainable population size” (https://www.fisheries.noaa.gov/feature-story/potential-biological-removal-levels-tool-conserve-marine-mammals). An analysis of the overlap between shipping activity and global range predictions of four whale species suggested that shipping occurs in 92% of these whales’ habitats. However, fewer than 7% of collision risk hotspots have management strategies in place to mitigate the risk of ship-whale collisions (*15*).

Mitigation measures are of three types (*16*): (1) routing measures to avoid sensitive or high-risk areas (e.g., (*17*)); (2) slowing down (e.g., (*18*)); and (3) avoidance manoeuvres after marine megafauna are detected, possible at small spatial and temporal scales (e.g., (*19*)).

Relatively small changes to ship behaviours, especially in key regions of the oceans, can have a big impact. For example, simulations show that decreasing vessel speeds by 10% across the global fleet could halve the risk of ships striking whales (*20*). For endangered North Atlantic right whales (*Eubalaena glacialis*), seasonally restricting vessels to speeds of 10kt or less was estimated to decrease whale mortality by 80-90% (*21*). After implementation of a 10kt speed restriction in 2008, expected seasonal mortality of North Atlantic right whales dropped by 22% (*22*).

Simultaneously, speed reductions can reduce greenhouse gas emissions, in line with the International Maritime Organization’s greenhouse gas (GHG) strategy that aims to reduce the shipping sector’s GHG emissions at least 50% below 2008 levels by 2050 (*23*). Baseline CO_2_ emissions for the global fleet from 2018-2030 could be reduced by 13, 24 and 33% with speed reductions of 10, 20 and 30% (*24*). Further, simulations show that small reductions in vessel speed can immediately and substantially reduce noise impacts on marine mammals (*25*).

However, avoidance of large areas and speed restrictions could disrupt world trade, and for the shipping industry there are concerns about the time and fuel costs of longer, slower routes, which might have significant economic costs; mitigation measures are unlikely to be voluntarily followed by shipowners if they increase the cost or time of delivering goods. Perhaps more realistic than a global reduction in fleet speed, is the use of traffic separation schemes that protect the local habitat (*26*). Making these adjustments is increasingly made possible through the availability of information on animal distributions and movements (e.g., (*13*)) alongside the improvements in voyage optimisation algorithms. Nevertheless, there is currently little understanding of the effect this might have on shipping routes, the additional cost, potential increase in CO_2_ emissions and the difficulty in managing these over seasonal megafauna movements that might result from these changes.

Here, we present a proof of concept demonstrating how information on the distribution of marine megafauna can be incorporated in voyage planning, using previously defined priority sites of the endangered whale shark (*Rhincodon typus*) as an example (*27*). We used a commercial Voyage Optimisation Software solution, T-VOS (Theyr Voyage Optimisation Solution), which incorporates the co-evolutionary Multi-Level Selection Genetic Algorithm with epigenetic blocking (cMLSGA) (*28*, *29*) to optimise 8 candidate vessel routes taking into account navigational restrictions, vessel performance, safety, meteorological and oceanographic conditions.

We demonstrate the voyage disruption that might occur for different scenarios, comparing the time and fuel increases that might occur from total avoidance of whale shark critical habitats or slowing down in these areas, and analysing the effects on a range of routes for different ship types.

## Results

### Benchmarking voyages optimised to protect biodiversity

When whale shark aggregation sites are treated as no-go areas, fuel loss is in the range of 0.12-7.21%, for the crude oil tanker (Figure 1a). These losses are inversely proportional to the distance added by avoiding the zone, as the vessel needs to increase its speed to recuperate the time lost when avoiding whale shark areas. This means that the longer voyages exhibit a relatively small percentage increase in fuel usage, <1.3%, but the shorter journeys exhibit a higher percentage increase. The exception is the Gulfport to Dorado voyage which has a low fuel loss, despite only making an approximately 2-day voyage. This is due to the adverse currents located directly within the core habitats, leading to base route deviating from the shortest route and thus partially avoiding the zone.

**Figure 1:**
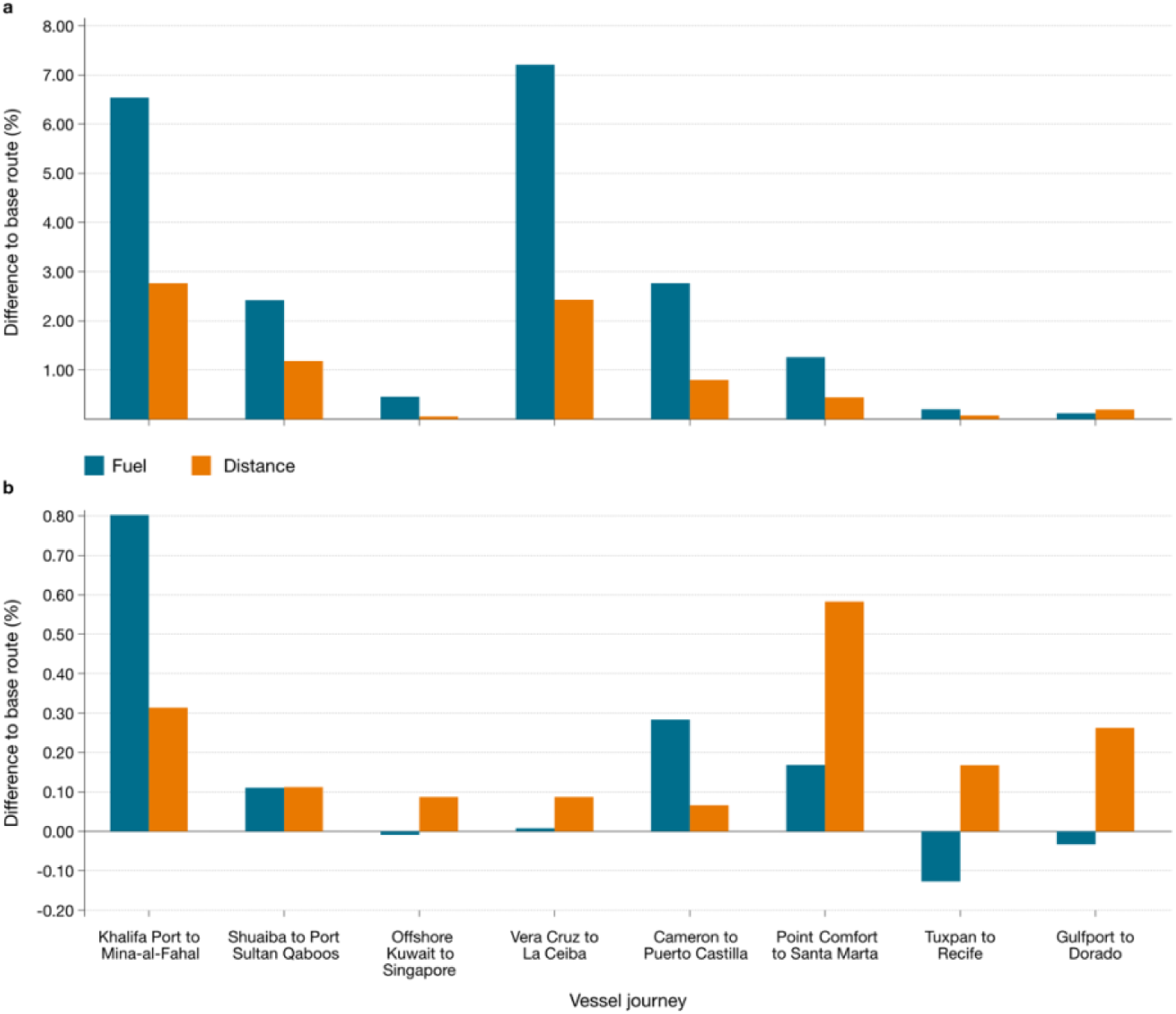
Impacts of whale shark ship-strike mitigation strategies on fuel-use and distance of simulated crude oil tanker voyages. Percentage change in fuel consumption (blue) and distance travelled (orange) for eight simulated crude oil tanker voyages when whale shark core habitats are treated as either (a) restricted navigation zones (no-go areas) or (b) speed reduction zones (10 kts limit), relative to baseline optimised routes. Rerouting (a) results in higher fuel use for shorter voyages, while longer voyages show minimal increases. Speed reduction (b) generally leads to lower fuel impacts due to limited speed adjustments and shorter travel paths.

Treating the whale shark habitats as speed reduction zones leads to a lower change in the fuel used, with Figure 1b showing minimal percentage changes of −0.13-0.8% in comparison to the base routes. This is because the vessel already travels at a low speed, 11-14kts, and the zones only require a minimal speed reduction. In addition, the vessels have shorter paths compared to re-routing and so the additional fuel used to make up for the loss of time due to slowing down in whale shark areas is minimal. For the shorter journeys, the speed reduction provides slight increases in fuel usage, changes of 0.01-0.8%. However, for the longer routes, the reduction in speed can be beneficial because the vessels are allowed to go slower than they could go due to party charter agreements. This means there is a feasibility that going through a speed reduction zone could cause a benefit, especially in charter party agreements that require the vessel to go at its maximum speed because it will allow the vessel to travel at a slower pace.

For the container ship, Figure 2 shows a similar behaviour to the oil tanker. The fuel loss is higher for the longer routes reaching up to 9.65%, a little higher than for the worst case in the oil tanker. The route with the highest fuel usage for the container vessel is the Khalifa Port to Mina-al-Fahal, rather than the Vera Cruz to La Ceiba route. However, for the longer voyages the impact on fuel consumption remains relatively small, <1%. There are also some slight differences in journey selection on the longer voyages, with the container vessel having negligible additional fuel usage on the Point Comfort to Santa Marta journey, while the oil tanker had a 1% increase in fuel usage. This is due to the oil tanker selecting a longer route compared to the base route. However, the Gulfport to Dorado route shows an increase of around 1.5% for the container ship, while this is negligible for the oil tanker, and in this case the additional distance travelled is the same.

**Figure 2:**
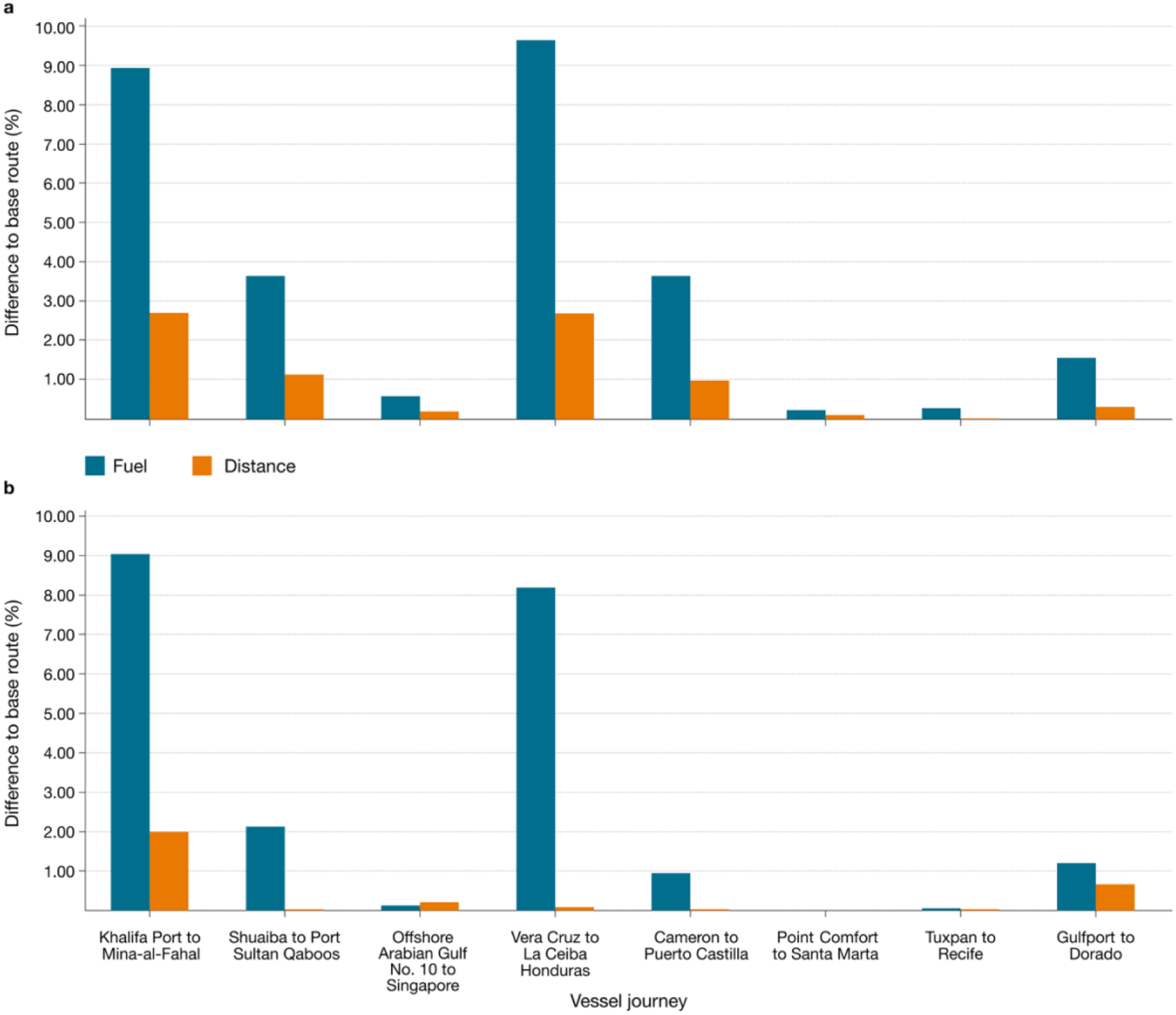
Impacts of whale shark ship-strike mitigation strategies on fuel-use and distance of simulated container ship voyages. Percentage change in fuel consumption (blue) and distance travelled (orange) for eight simulated container ship voyages when whale shark core habitats are treated as either (a) restricted navigation zones (no-go areas) or (b) speed reduction zones (10 kts limit), relative to baseline optimised routes. Fuel increases are greater for shorter voyages, where added distance has a proportionally larger impact. Longer voyages show smaller relative increases in fuel use due to greater opportunity for route flexibility and speed optimisation.

The speed reduction zone is more detrimental to carbon reduction for the container ship (Figure 2b). The change in fuel for the shorter routes remains high, with values from 3.9%-9.1% which is a similar increase to the no-go routes. The longer routes again show a minimal increase with values of 0%-1.1%; in these cases, the change in the distance being travelled is minimal.

To illustrate the change in routes between the two types of vessels with distinct speed profiles, the Khalifa Port to Mina-al-Fahal voyage is shown on Figure 3 with the pink line representing the oil tanker and the purple line representing the container route. In this scenario the oil tanker follows a path that is similar to the shortest path, where the vessel enters the habitat quite early and further south, making a minimal effort to avoid the area, focusing on reducing the distance to the goal and avoiding directly sailing into the stronger currents. The container vessel enters the whale shark area at the most northern part, skirting around the zone before using a gap in the habitat that allows it to minimise the time spent in the speed reduction zone. This alleviates the time spent with the more severe speed reduction but requires sailing directly into strong currents. This increases the distance travelled and has a more severe penalty to the fuel usage.

**Figure 3.**
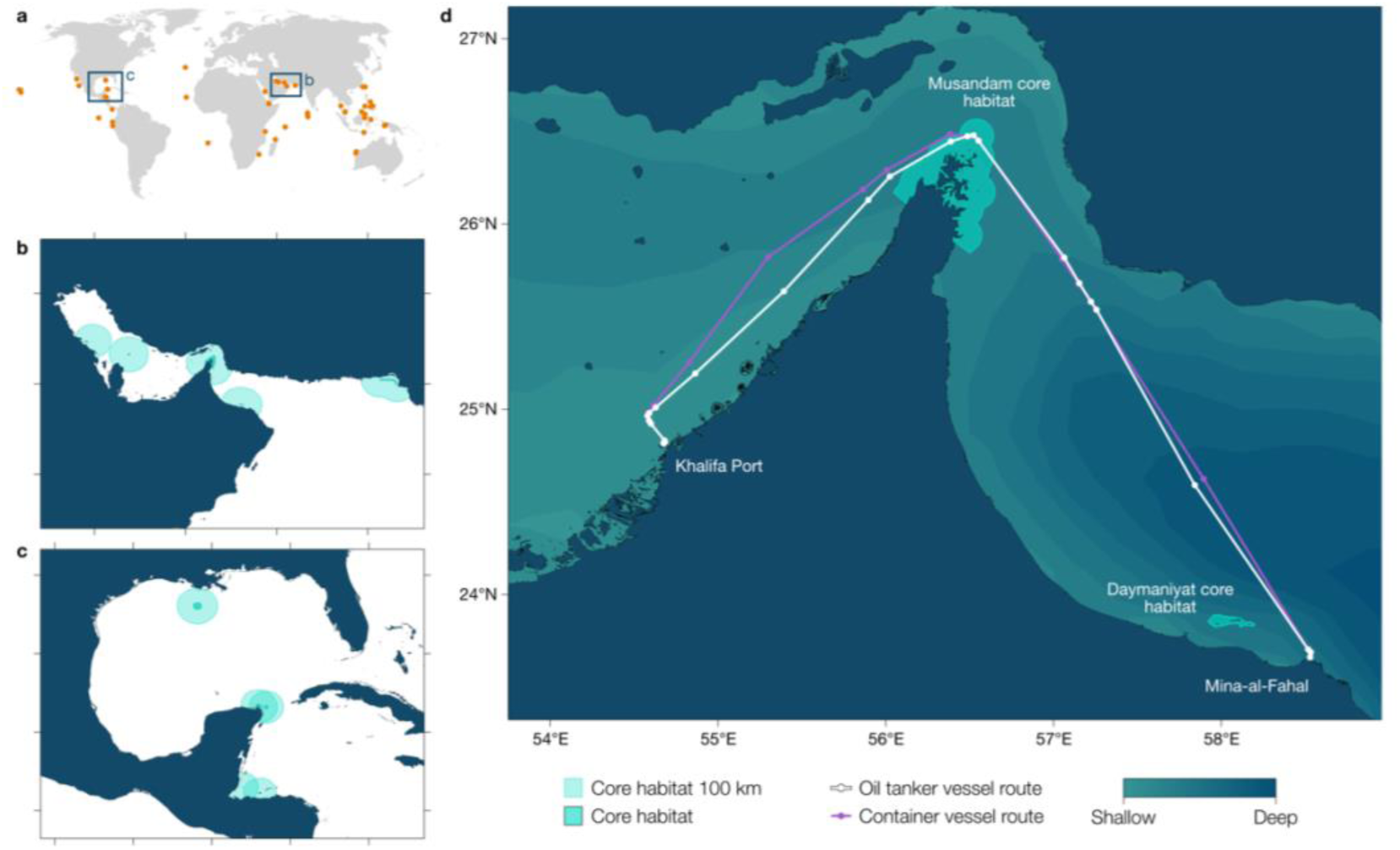
Simulating speed reductions in global whale shark core habitats. Maps of whale shark aggregation sites shown at (a) global scale, (b) in the Persian Gulf, and (c) in the Gulf of Mexico. Core habitat polygons (dark shading) represent areas with the highest density of whale shark observations, compiled from decades of monitoring and expert input. Panel (d) shows optimised routes for a crude oil tanker (white) and container vessel (purple) between Khalifa Port and Mina-al-Fahal, one of eight candidate voyages used in simulations, intersecting the Musandam whale shark aggregation hotspot. In this scenario, core habitats are treated as speed reduction zones with a maximum vessel speed of 10 kts.

## Discussion

Targeting management of damaging human activities within key sites for marine megafauna is now essential for their conservation (*30*). Management strategies that consider both the ecological needs of at-risk species and the potential operational impacts to industry are needed to facilitate swift uptake of regulations, as well as long term stakeholder adherence (*31*). Here we show that by incorporating the location of priority habitats for whale sharks directly into ship voyage optimisation software during the planning phase, important areas can be protected while having negligible impact on overall shipping fuel costs. We find that mitigation could be tailored to the type of ship voyage to minimise industry disruption, with success hinging on proper implementation and adherence.

For longer voyages, over 6 days, full avoidance of whale shark priority areas is recommended with no severe adjustments to vessel’s speed required, as at most 1% extra fuel will be consumed at the lower end of the boundary and less than 0.5% extra on voyages lasting more than 12 days. In contrast, for shorter voyages, a reduction in speed is recommended rather than avoiding whale shark priority areas, as it can limit potential fuel losses to less than 1% for slower vessels, whereas a complete avoidance of speed reduction would significantly increase operational costs.

Many vessels operate under charter party agreements that strictly define permissible speed ranges. These vessels are typically not permitted to reduce their speed unless explicitly allowed by the terms of the agreement or by specific legislation that permits such actions, indicating the need for legislative amendments and the role of organizations such as the International Maritime Organization (*20*, *26*). Here, biodiversity priority zones for at-risk species could be built into charter party agreements, relying on evidence-based spatial layers to be generated and passed to the correct parties.

For high-speed vessels with restrictive operational ranges, such as larger container ships or liquefied natural gas carriers, speed reduction may cause more problems than rerouting, due to engine constraints and the need to switch to auxiliary engines.

Furthermore, it will not prevent significant fuel increases when whale shark priority areas are being considered on short-duration voyages. However, these vessels rarely operate solely on such scenarios. Instead, shorter routes are typically integrated into longer voyage plans involving a series of interconnected passages and port calls. Therefore, fuel losses can potentially be mitigated by optimising port call timings and slightly increasing the speed on subsequent passages, although this would require a more significant change to charter party agreements. Voyage optimisation solutions can already achieve the necessary optimisation to the route. More broadly, these techniques are not unique to shipping, and there are parallels that could be drawn in the aerospace industry and potentially other transport systems to improve biodiversity.

Traffic separation schemes already provide some protection for marine megafauna in critical areas (*32*). However, it can be a slow process to introduce new schemes. Some schemes are static throughout the year, while other schemes operate seasonally(*32*). Many megafauna will migrate through the year and protected areas or traffic separation schemes need to be dynamic to protect ocean biodiversity (*33*). For example, whale sharks are known to aggregate within the priority sites included in our voyage simulations on a seasonal basis, meaning that mitigation need only operate within specific time windows.

This would further reduce impacts to operations. However, with climate change the habitats of megafauna (*34*, *35*) and optimal routes for vessels could change more often, making adaptation and resilience to this change an important factor for any proposed solution. The current technology can cope with regular updates to schemes, and this would seem to provide minimal impact to the fuel use of most journeys.

### Priorities for optimizing ship voyages to mitigate ship strike

Here we highlight the importance of considering journey length when planning for marine megafauna ship collision mitigation to ensure global shipping itself is not impacted too heavily. We also demonstrate the importance science-industry engagement through providing evidence-based data on priority marine megafauna areas in known high risk zones as per the suggestions outlined in Womersley et al. (*31*). At present, a reduction in the total number of ships moving through these priority habitats provides the best solution to mitigate shipping impacts, but having some vessels moving slowly (i.e., <100% compliance on avoidance) will still provide benefits to biodiversity. Furthermore, it is essential to consider the whale shark areas dynamically, as static databases currently do not consider species’ seasonal movements or potential relocations in future due to climate change.

Building on these recommendations, we present four priorities for mitigating ship strike through biodiversity prediction and voyage optimisation.

#### Improved data and processing pipelines to generate accurate and precise predictions

Improved data and processing pipelines are essential for generating accurate and precise priority areas to integrate into vessel voyage optimisation software. Basic distribution data are lacking for many marine megafauna species. Even where data do exist, they may not be FAIR (findable, accessible, interoperable, reusable), and/or too sparse for accurate predictions. Nonetheless, biodiversity datasets collected through human observations and various sensor observations are rapidly growing (*36*), including data on the distribution of marine megafauna (e.g. (*37*, *38*)). For example, fast-growing repositories of biologging (animal tracking) data provide data on the locations of tagged animals at varying time lags (*39*) and can be used to predict the probability of animals being in a given location. Static or glider-mounted networks of passive acoustic recorders could be used for real-time detection of vocalising marine megafauna such as whales and dolphins (*40*, *41*). Data integration methods can be used to combine different types of location information (e.g., (*15*, *42*, *43*)). Nonetheless, latency--the time between data collection and prediction--can be long: often data are years old by the time they are used to predict the distribution of marine megafauna (e.g., (*44*)). Enhancing the speed of biodiversity data processing and ensuring timely dissemination to relevant stakeholders can significantly improve decision-making, but also relies on science-industry communications. To facilitate this, in the first instance designated priority areas should be small and precise to promote greater adoption by minimising disruption to shipping. It is important to prevent the implementation of overly broad geographical restrictions, which can increase operational costs without providing substantial benefits to megafauna conservation. For example, shipping occurs in 91.5% of the predicted distribution of blue, fin, humpback and sperm whales, but ship-strike risk ‘hotspots’ (areas with the highest coincidence of shipping and predicted whale distribution) could all receive spatial protection by expanding current speed-reduction zones by 2.6% of the ocean’s surface (*15*).

#### Dynamic, real-time adaptability and forecasting capabilities

The marine realm is highly dynamic, and building systems with dynamic, near real-time spatiotemporal adaptability and forecasting capabilities at various management-relevant timescales is vital for effectively managing species protection while minimising disruptions to shipping operations. Moving forward, incorporating dynamic priority areas, which consider both space and time, will allow for better tracking of predictable species movement/distribution (e.g. annual migrations, seasonal aggregations) and potential redistributions (e.g. poleward shifts) caused by climate change, while helping to minimise impacts on shipping. It is critical that forecasts are iteratively improved, including new data (and new data types) as these become available, serving to reduce uncertainty in predictions and ensuring that spatial data remains reliable over time and useful for operational decision-making (*45*, *46*). Forecasts also need to be able to deal with climatic extremes (*47*). To achieve a responsive and resilient system, industry partners need to be updated when new information on priority site location or timing arises. Whale Safe (https://whalesafe.com), for example, is a mapping and analysis tool that combines predictions of whale distribution (*48*) with real-time presence data streams (*49*) to provide a daily ‘whale presence rating’ for the Santa Barbara Channel and the San Franciso region to the shipping industry, natural resource managers and the public through the Whale Safe website. Globally, such forecasts might be served through organisations such as the IMO, and we have demonstrated here how these information layers can then be incorporated by commercial voyage optimisation software.

#### Regulatory and incentive-based compliance while monitoring effectiveness

Effective regulatory frameworks and incentive-based compliance mechanisms are essential to balance shipping operations with the protection of marine megafauna. Collaboration with the International Maritime Organization (IMO) and policymakers will be key to implementing speed reduction or rerouting measures in sensitive habitats. Given that many vessels operate under charter party agreements that restrict speed changes, contractual flexibility will be necessary to permit adjustments in ecologically critical areas, such as priority whale shark habitats.

Incentive-based approaches could complement regulation, for example through ‘badging’ or eco-labelling compliant voyages, as has been achieved in other sectors (*50*). Rebates on port dues for ships travelling slowly (*51*) or targeted fuel levies (*52*) could also encourage adherence. The IMO’s April 2025 approval of a Net-zero Framework (https://www.imo.org/en/mediacentre/pressbriefings/pages/imo-approves-netzero-regulations.aspx) offers an additional model for integrating conservation incentives. This framework introduces a global fuel standard for greenhouse gas emissions, requiring vessels that exceed Greenhouse Fuel Index (GFI) thresholds to compensate for surplus emissions through credit transfers, banked units, or offset purchases. Ships operating with zero- or near-zero-emission technologies qualify for financial rewards, funded through a dedicated Net-zero Fund that will support research, development, and deployment of low-carbon maritime technologies. Similar mechanisms could be adapted to biodiversity protection, linking compliance with speed reductions or rerouting in priority habitats to existing emissions-based incentives—thereby aligning climate and conservation objectives while promoting industry uptake.

Assessing the conservation benefits of route adjustments, implementing systems to track both compliance and ecological outcomes, and conducting realistic cost simulations of mitigation strategies will further help optimise decision-making. Addressing data ownership, clarifying responsibilities for forecast updates—particularly in Areas Beyond National Jurisdiction—and ensuring adequate resources for generating and updating spatial predictions will be critical. These steps will also support the effective integration of biodiversity data into maritime routing processes.

### Integration with existing maritime navigation and safety systems with a focus on multiple species where possible

Integrating conservation measures with existing shipping navigation frameworks and safety systems is crucial for protecting marine megafauna from collisions while maintaining operational efficiency. For example, expanding and adapting traffic separation schemes to account for priority areas such as whale shark aggregations can enhance both navigational safety and species protection. Incorporating spatial habitat data, such as through initiatives like WhaleSafe into these systems allows for more targeted and effective routing decisions. However, it is important to avoid overly broad avoidance zones, as they may undermine stakeholder trust and can significantly raise operational costs. This will be particularly challenging in areas used by multiple species, as each has varying levels of exposure, vulnerability, and ecological significance. In these cases, developing risk-weighted models that account for these differences, including lesser-known or data-deficient species, can help ensure that conservation efforts are appropriately targeted and inclusive. This relies on robust data and effective dissemination in areas used by multiple species. Where data dissemination is challenging, exploring the use of umbrella species, whose protection can indirectly benefit a wide range of other organisms sharing the same habitat, might offer a promising strategy for maximizing ecological impact with limited resources.

## Conclusions

We demonstrate that spatial information on marine megafauna habitats can be directly incorporated into commercial voyage optimisation, enabling the shipping industry to evaluate the operational implications of two common ship-strike mitigation strategies: speed reduction and area avoidance. Using whale shark priority sites as a proof of concept, we show that mitigation can be tailored to voyage length and vessel type, with speed reduction most suitable for shorter routes and area avoidance for longer voyages. In both cases, fuel and time penalties were minimal, suggesting that biodiversity considerations can be integrated into routing decisions without substantial disruption to global trade.

Realising this potential at scale will require coordinated action across four areas. First, improved data pipelines are needed to generate accurate, high-resolution priority areas for a broader range of species. Second, dynamic, near real-time forecasting capabilities must be developed to capture seasonal movements and long-term shifts in species distributions. Third, regulatory and incentive frameworks—such as those recently introduced under the IMO’s Net-Zero Framework—should be adapted to incorporate biodiversity protection, aligning conservation goals with decarbonisation targets. Finally, integration of biodiversity layers into existing maritime navigation and safety systems should be pursued, with a focus on multi-species management and risk-weighted protection measures.

By embedding biodiversity into voyage optimisation, the maritime sector can contribute meaningfully to global efforts to reduce ship strikes, safeguard at-risk species, and maintain the resilience of ocean ecosystems, while sustaining the efficient movement of goods on which the global economy depends.

## Methods

Novel, and realistic, time and fuel implications are provided for the disruption to eight candidate routes, that currently run through the habitats of whale sharks. A comparison is made to current shipping, a reduction in speed to 10kts and rerouting the vessels around the habitats entirely. Two types of vessels are compared, a slower oil tanker and a more rapid container ship.

### Voyage optimisation implementation

T-VOS (Theyr Voyage Optimisation Solution) incorporates the co-evolutionary Multi-Level Selection Genetic Algorithm with epigenetic blocking (cMLSGA) (*28*, *29*). This algorithm is inspired by principles of evolution from the extended synthesis, epigenetics and multilevel selection. In Multi-Level Selection the survival of the individuals is not solely determined by their own fitness but also that of the collective that they belong to, similarly to species with hierarchical or pack-based structures. cMLSGA has demonstrated strong performance in addressing complex engineering problems compared to other genetic algorithms and has been demonstrated to have leading performance on Voyage Optimisation problems (*9*). This approach is now extended with Epigenetic blocking mechanisms that have been shown to provide excellent performance on dynamic multi-objective problems (*28*). To mimic epigenetic mechanisms, variables in each individual are probabilistically blocked during crossover. After crossover, genes chosen to be blocked in the offspring are set to the unblocked parent’s genes. Any mutation steps occur after the epigenetic process to prevent overwriting changes made during mutation.

The multi-objective facilitates the simultaneous optimization of conflicting objectives. This is crucial on complex multi-objective problems that involve numerous variables and constraints like the voyage optimisation where at least two main objectives are present, usually fuel consumption and voyage duration.

The hyperparameters used in this paper are the standard values used in T-VOS, which were refined based on comprehensive benchmark on actual voyages and remain proprietary. The total number of function calls is limited to 500,000. This leads to a total optimisation time of less than one minute, for voyages under 60 days, and <0.2% effectiveness loss in comparison to tests with 3,000,000 function calls indicating convergence.

### Whale shark aggregation sites

Whale sharks have long been known to be at risk of ship strike with some of the first records of collisions documented in the early 1900s (*53*), but only recently has the issue gained traction due to partially unexplained population declines of >50% in 75 years(*54*). Whale sharks spend almost half of their time in surface waters – within the range of passing ship hulls – and in general show limited avoidance to vessels (*13*). The species ranges widely, crossing ocean basins and travelling upwards of 10,000km annually, but also predictably aggregates in large numbers at approximately 50 sites around the world (*27*). Many of these sites are small, defined areas driven by sea surface temperature and ephemeral prey availability(*55*). They primarily consist of juveniles (with a bias toward males) slowly feeding at the surface. Aggregations have recently been put forward as the most tractable way to meet conservation targets for whale sharks by focusing mitigation efforts in discrete, predicable locations (*56*). There is currently no mitigation to limit strikes to whale sharks anywhere globally despite a new resolution aimed at tackling this issue being adopted at the Convention on the Conservation of Migratory Species of Wild Animals 14^th^ Conference of Parties in early 2024 (*57*). Therefore, although the species has been shown to be at risk from ship strike across large portions of its range (*13*), trialling mitigation strategies within identified priority aggregation sites provides an opportunity to protect areas where the most individuals gather whilst safeguarding key demographics (*27*).

Through compiling spatial information provided by local researchers working at each aggregation site, a recent study mapped all known core habitats and buffer zones – that is, where the most individual whale sharks gather and anywhere locally where they have been seen, respectively – based on decades of monitoring (*27*). In the study, sites were ranked based on relative levels of local shipping activity, revealing those situated in the Persian Gulf and the Gulf of Mexico to be among the most dangerous for whale sharks. These regions were selected as case studies to integrate into the T-VOS software, and 11 aggregation site polygons for core habitats and buffer zones were made available for analysis.

### Optimising voyages accounting for whale shark aggregation sites

To investigate the impact of the whale shark habitats on the routing, the aggregation site polygons for core habitats and buffer zones were directly integrated into the T-VOS software. These are considered as speed reduction zones, with speed limited to 10kts, or restricted navigation areas where no sailing is allowed. Eight port-to-port voyages are simulated, selected based on their proximity to the whale shark habitats while providing a range of different lengths of voyage between major ports where larger vessels are likely to operate (Table 1).

**Table 1.** Summary of simulated port-to-port voyages used in routing optimisation. Origin, destination, and approximate distance of eight commercial shipping routes selected for voyage optimisation simulations. Route originating in the Persian Gulf (a) and Gulf of Mexico (b) were chosen based on proximity to whale shark aggregation sites and represent a range of voyage lengths and geographic contexts. These were used to evaluate the impact of speed reduction and area avoidance strategies on fuel consumption and distance travelled.

| Origin | Destination | Approximate distance (km) |
| --- | --- | --- |
| *a) Persian Gulf* |  |  |
| Khalifa Port (UAE) | Mina-al-Fahal (Oman) | 672 |
| Shuaiba (Kuwait) | Port Sultan Qaboos (Oman) | 1,325 |
| Offshore Kuwait | Singapore | 7,157 |
| *b) Gulf of Mexico* |  |  |
| Vera Cruz (Mexico) | La Ceiba (Honduras) | 1,731 |
| Cameron (USA) | Puerto Castilla (Honduras) | 1,953 |
| Point Comfort (USA) | Santa Marta (Colombia) | 3,075 |
| Tuxpan (Mexico) | Recife (Brazil) | 7,918 |
| Gulfport (USA) | Dorado (Mexico) | 1,161 |

The current traffic separation scheme in the Persian Gulf is ignored, this is because it currently takes vessels through the whale shark area. This means the simulations are equivalent to having a speed limit of 10kts in the traffic separation scheme zone or moving the zone north, in the case of the no-go simulation.

All voyages are simulated for a crude oil tanker with an approximate length of 336m and the deadweight tonnage of 300,000t and container ship of 366m and 133,600t respectively, both under laden conditions. The models use specific fuel-oil consumption curve (SFOC), based on real operational data, as a baseline for the fuel consumption and the Townsin and Kwon empirical method for the added resistance due to met-ocean conditions. The operational speed ranges are 11-14kts, based on the average vessel speed of 12.5kts, for the tanker and 18-21kts with 19.5kts average for the container vessel, to simulate the conditions where the vessel must slow-down in the speed reduction zones.

During the optimisation, the vessel is allowed to change its course every 55 nautical miles (NM) and its speed every 275NM, and each scenario is simulated three times to provide statistically significant results. The width of optimisation mesh is set to 1000km, which defines the maximum perpendicular distance the vessel is allowed to navigate away from the shortest great circle route during the optimisation. For each case, the best on-time fuel optimised route, defined as the route with arrival at port no later than the contractual deadline, is evaluated and compared against the base route. The base routes are defined as optimised voyages without taking the whale shark areas into account and with the traffic separation schemes ignored. The latest allowed arrival time is based on the voyage time with average speed of 12.5kts. All voyages start on the same date, 1st October 2022 10:00 UTC, to minimise the influence from met-ocean conditions and remove the uncertainty bound to operating on data forecasts.

## Author contributions

R.R.R: Conceptualization; investigation; writing – original draft; writing – review and editing. P.A.G.: Conceptualization; data curation; formal analysis; investigation; methodology; software; writing – original draft; writing – review and editing. F.C.W.: Data curation; formal analysis; investigation; visualization; writing – original draft; writing – review and editing. D.W.S.: Investigation; writing – original draft; writing – review and editing. A.J.S.: Conceptualization; investigation; methodology; software; writing – original draft; writing – review and editing.

## Acknowledgements

We would like to thank the whale shark researchers listed as authors on the publication “Identifying priority sites for whale shark ship collision management globally” published in Science of The Total Environment (https://doi.org/10.1016/j.scitotenv.2024.172776) who helped to delineate the spatial whale shark habitat data used here. F.C.W was supported by the Marine Research and Conservation Foundation (MARECO) while compiling the whale shark habitat data and received additional support from a UK Natural Environment Research Council Southampton INSPIRE DTP Studentship (no. NE/S007210/1). F.C.W and D.W.S. were supported by a European Research Council Advanced Grant (no. 883583 OCEAN DEOXYFISH) awarded to D.W.S. within the EU Horizon 2020 Programme. A.J.S. was supported by Lloyd’s Register Foundation as part of the Sustainability mission in the Alan Turing Institute.

## Declaration of generative AI and AI-assisted technologies in the manuscript preparation process

During the preparation of this work the author(s) used ChatGPT and Perplexity in order to assist with text editing and literature review. After using this tool/service, the author(s) reviewed and edited the content as needed and take(s) full responsibility for the content of the published article.

